# Locally restricted glucose availability in the embryonic hypocotyl determines seed germination under ABA treatment

**DOI:** 10.1101/2021.04.07.438879

**Authors:** Xueyi Xue, Ya-Chi Yu, Yue Wu, Huiling Xue, Li-Qing Chen

## Abstract

- Abiotic stresses directly affect seed germination, plant growth, and reproduction. Seed germination is a complex and energy-demanding process, involving diverse physical, metabolic and cellular events, and is controlled by intrinsic and environmental cues. Abiotic stresses involving abscisic acid (ABA) inhibit seed germination by suppressing physiological processes essential for completion of germination. It remains unclear whether the component tissues of an embryo respond to stress, and are suppressed differentially.
- Here we used a genetically engineered FRET sensor to show that ABA significantly affected the spatiotemporal distribution of glucose (Glc) in lower hypocotyl region. Transcriptome analysis and ^14^C-Glc uptake assay suggested that Glc limitation in the embryonic hypocotyl was largely due to suppressed sugar partitioning along with enhanced sugar metabolism. The loss-of-function mutants of ABA-induced sucrose-phosphate synthase (*SPS*) genes accumulated more Glc, leading to ABA insensitivity during germination. In addition, we identified molecular signatures that Glc antagonizes ABA by globally counteracting the ABA influence on gene expression, including expansin (*EXP*) family genes that are inhibited by ABA.
- This study presents a new perspective on the interaction between ABA and Glc in response to external stimuli, which restricts Glc availability and thereby controls the transition from seed to seedling.

## Introduction

In addition to global warm, climate changes have been leading to more frequently severe abiotic stresses on plants, such as heat waves, droughts and salinity, which can directly affect food production, due to the destructive effects on seed germination, plant development and reproductive growth (Barnabas *et al*., 2008; Raza *et al*., 2019). Seed germination is a complex process including diverse physical, metabolic and cellular events, such as recovery from the drying, resumption of basal metabolism, DNA repair and mitochondria rebuilt (Bewley, 1997). Germination starts with the water uptake by the dry seeds and terminates with emergence of root radical (Bewley, 1997), which is often the first visible sign of germination in model plant Arabidopsis thaliana. Germination has been recognized to result from cell elongation, but not cell division, in lower hypocotyl region to push the radicle out (Sliwinska *et al*., 2009).

An *Arabidopsis* seed consists of the symplasmically isolated seed coat, endosperm and embryo, the latter including four symplasmically isolated components: the cotyledon, shoot apex, hypocotyl, and root (Kim & Zambryski, 2005). The cell-to-cell communication among these sub-domains is limited via plasmodesmata, while it can be facilitated by protein carriers on the plasma membrane (PM). The cotyledons of embryos are differentiated into the major storage organs for lipids, protein, and sugars (Baud *et al*., 2008), while the cell elongation takes place in the hypocotyl during germination. Cell elongation is an energy-consuming process, associated with cell walls loosening and remodeling (Cosgrove, 1999; Refregier *et al*., 2004). Sugars derived from storage remobilization serve as a source of energy and nutrients for cell wall biosynthesis (Reiter, 2002; Verbancic *et al*., 2018). It remains unclear if sugar, derived from seed reserves in the hypocotyl, is sufficient to fuel the cell elongation to finish seed germination or if subsequent sugar transporters mediated sugar partitioning among the subdomains can drive sufficient hypocotyl elongation for germination to occur under stresses.

Several sugar transporter families have been identified in Arabidopsis so far. Sugar transport proteins (STPs) are symporters to transport monosaccharides with H^+^ (Boorer *et al*., 1994; Sherson *et al*., 2000). Sucrose transporters (SUCs) are energy-dependent H^+^/sucrose symporters (Sauer & Stolz, 1994; Gottwald *et al*., 2000). The tonoplast monosaccharide transporters (TMTs) functions as H^+^/monosaccharide antiporters at vacuole membrane (Wormit *et al*., 2006; Wingenter *et al*., 2010). The sugars will eventually be exported transporters (SWEETs) are uniporters that facilitate sugar diffusion across membranes along the gradient (Chen *et al*., 2010; Chen *et al*., 2012). But it is not clear whether these sugar transporters are involved in the sugar-ABA interaction.

Förster resonance energy transfer (FRET)-based sensors have been used to monitor the steady state substrate levels with different affinities in mammalian cells (Fehr *et al*., 2003; Lager *et al*., 2003; Okumoto *et al*., 2005). Several genetically modified FRET Glc nanosensors have been used to assess Glc levels between different cells and tissues with or without treatments in Arabidopsis (Deuschle *et al*., 2006). This noninvasive imaging technique has been explored to investigate cellular dynamics of ABA, gibberellin, MgATP^2-^ and Ca^2+^ in plants (Krebs *et al*., 2012; Jones *et al*., 2014; De Col *et al*., 2017; Rizza *et al*., 2017). In this study, we found that in the presence of abiotic stresses and ABA, Glc uptake in germinating seeds was suppressed. FRET sensor shown that Glc availability in embryonic hypocotyl region is limited upon ABA treatment. Transcriptomic analysis uncovered that the gluconeogenesis slows down and Glc supply was locally restricted resulting in a deficiency of sugar in the embryonic hypocotyl under ABA treatment, impairing the elongation required for the completion of seed germination. This study provides a new perspective on the antagonistic effects of ABA and sugar on germination.

## Materials and Methods

### Plant material and growth conditions

*Arabidopsis thaliana* wild-type plants are Columbia background (Col-0). Seeds of FLIPglu-170nΔ13 and FLIPglu-3.2mΔ13 in *rdr6* background were obtained from Wolf B Frommer (Deuschle *et al*., 2006). The CaMV 35S promoter of pPZP312 (digested by XhoI and ApaI) was replaced by a fragment containing the UBQ10 promoter and a kanamycin resistance cassette flanked by SalI and BstXI sites (XhoI and ApaI compatible ends, respectively), simultaneously introducing PstI and BamHI to flank the UBQ10 promoter. The *sps1,4* and *sps1,2,3* mutants were obtained from Javier Pozueta-Romero (Bahaji *et al*., 2015). Seeds were sterilized by 70% EtOH for 5 minutes, rinsed three times with distilled water, and then imbibed at 4 °C for 3 days. Seeds were sown on ½ MS medium with 1% agar (A1296, Sigma), pH 5.8. Sugars, salt, and hormone were sterilized by 0.22 μm filter. Plates were incubated in growth chamber with constant light (100 µmol/m^-2^/s^-1^) at 22 °C.

### Glc uptake assay in germinating seeds

After 3 days imbibition, seeds were sown on ½ MS media with 1% agar and grown under constant light, 22 °C for 24 hours, then transferred to fresh liquid ½ MS medium supplemented with ^14^C-Glucose (NEC-042X, PerkinElmer) and a stress treatment for 6 hours with shaking at 200 rpm. Stress treatments were ABA (AC133480010, Fisher Scientific), sodium chloride (BP-358-212, Fisher Bioreagents), mannitol (M4125, Sigma) or heat treatment at 38 °C in a water bath. Seeds were dissected to embryo and seed coat (containing endosperm and testa). Twenty seeds/embryos per replicate were transferred into a 1.5-ml tube and ground by pestle, followed by adding 200 µL 1% bleach. Radioactivity was measured by liquid scintillation spectrometry for 3 minutes per vial.

### Sugar measurement

After 3 day of imbibition, seeds were incubated in liquid ½ MS medium with or without treatment under constant light, 22 °C for 24 hours. Seeds were homogenized by pestle in a 1.5-ml tube. Total soluble sugars were extracted with 80% ethanol. After the ethanol was evaporated, and the sugars were dissolved in water, the sugar content was measured by high performance liquid chromatography (HPLC).

### Fluorescence microscopy

Seeds were dissected right before imaging and keep in corresponding ½ MS liquid medium. Fluorescence imaging of sensor embryos was performed on an inverted epifluorescence microscope (Carl Zeiss LSM710). Confocal images were acquired using Plan-Apochromat 20x/0.8 M27 objective. The 458- and 514-nm lasers were used for excitation of donor CFP and acceptor YFP. Fluorescence emission was detected by PMT detectors at 461-500 nm for donor and 525-575 nm for acceptor. Images were taken simultaneously with FRET donor and acceptor emission under donor excitation (458 nm), while acceptor emission under acceptor excitation (514 nm).

### Image processing and analysis

Image processing and analysis were performed using FIJI (Schindelin *et al*., 2012). The ratio image was processed and analyzed as described (Jones *et al*., 2014). Briefly, the ratio in the entire embryo was calculated by summing the Z-stack slices (interval = 10 µm) of acceptor FRET_YFP and donor FRET_CFP were summed, and then subtracting the YFP channel image by automatic local thresholding (method = Bernsen radius = 15 parameter_1 = 0 parameter_2 = 0 white). DxDm and DxAm intensities were filtered by Gaussian Blur; sigma = 1 scaled. The DxAm/DxDm ratio was calculated for each image, multiplied by the image of the YFP channel, and then divided by 255. The calculated images were displayed in 16 colors and the brightness/contrast was adjusted to 1-3.8.

### Seed sampling and mRNA-seq library preparation

After 3 day of imbibition at 4 °C, WT seeds were sown on ½ MS medium and incubated in the growth chamber under constant light, 22 °C for 24 hours. Seeds were transferred to ½ MS liquid medium supplemented with either 60 mM Glc, 5 µM ABA or both for 6 hours. Four biological replicates were collected for each treatment. Total RNA was extracted by RNeasy PowerPlant Kit (13500-50, Qiagen), DNA was removed via Turbo DNA-free kit (AM1907, Invitrogen). The cDNA libraries were prepared from 1000 ng RNA using KAPA mRNA Hyper Prep kit (KK8581, Roche) with KAPA Dual-indexed Adapter kit for Illumina platforms (KK8722, Roche). The libraries were sequenced by NovaSeq 6000 (Illumina) with paired-end reads at Roy J. Carver Biotechnology Center, UIUC.

### Transcriptome analysis

Gene expression level was quantified using Kallisto v 0.44.0 (Bray *et al*., 2016) by mapping to *Arabidopsis thaliana* (version 11) primary transcript sequences (https://phytozome.jgi.doe.gov). Then differentially expressed genes (DEGs) were identified using Sleuth (Pimentel *et al*., 2017) program and were those showing |b| ≥ 0.2 and *q*-value < 0.05 with expression greater than 10 counts per million reads in at least 50% samples. Samples were evaluated based on pair-wise comparison between different treatment conditions and outliers within each treatment condition were removed for the further analysis based on principal component analysis results. Hierarchical clustering was performed using gplots package in R based on the TPM data for DEGs. GO analysis was performed using PANTHER (Mi *et al*., 2019) with significant *P* values (*P* < 0.05) by filtering out the top hierarchical GO terms. Pathways with highlighted genes were downloaded from DAVID Bioinformatics Resources 6.8 (Huang da *et al*., 2009).

### Statistical analyses and data presentation

All statistical analyses were performed using either GraphPad Prism v.9.0.0, OriginPro v.9.55 or excel. Two tailed *t*-test was used for comparisons between two groups. For the graphs that did not show the individual data points, the individual values were listed in the file of source data.

## Results

### Abiotic stresses and ABA suppress Glc uptake in germinating seeds

Here we examined seed germination under salinity, osmosis, heat, and ABA stress. Consistent with previous studies (Pratap & Sharma, 2010; Vallejo *et al*., 2010; Chiu *et al*., 2012; Wilson *et al*., 2014), these stresses dramatically delayed seed germination (Fig. 1A), with the greatest inhibition by ABA treatment. As abiotic stress inhibits many cellular events essential for completion of germination (Daszkowska-Golec, 2011), we examined the impacts of stresses including ABA on sugar transport in germinating seeds. Quantified radiotracer uptake showed that the stresses suppressed Glc uptake (Fig. 1B and Fig. S1) and that ABA inhibition of Glc uptake was dosage-dependent (Fig. 1C). Since ABA-dependent signaling plays critical roles in stress response (Seo *et al*., 2000; Xiong *et al*., 2001) and the Glc uptake inhibition by stresses was similar to that triggered by ABA (Fig. 1A and 1B), we focused only on ABA treatments for the remaining experiments. Unsurprisingly, Glc, sucrose and fructose accumulation were all reduced by ABA (Fig. 1D), implying that ABA may restrict both sugar allocation and metabolism during germination. This notion was supported when addition of Glc largely rescued the germination delay triggered by ABA (Fig. 1E), consistent with previous studies (Finkelstein & Lynch, 2000). We also determined that 60 mM Glc was the most effective, although 15 mM was sufficient to antagonize ABA inhibition during seed germination (Fig. S2).

**Fig. 1.**
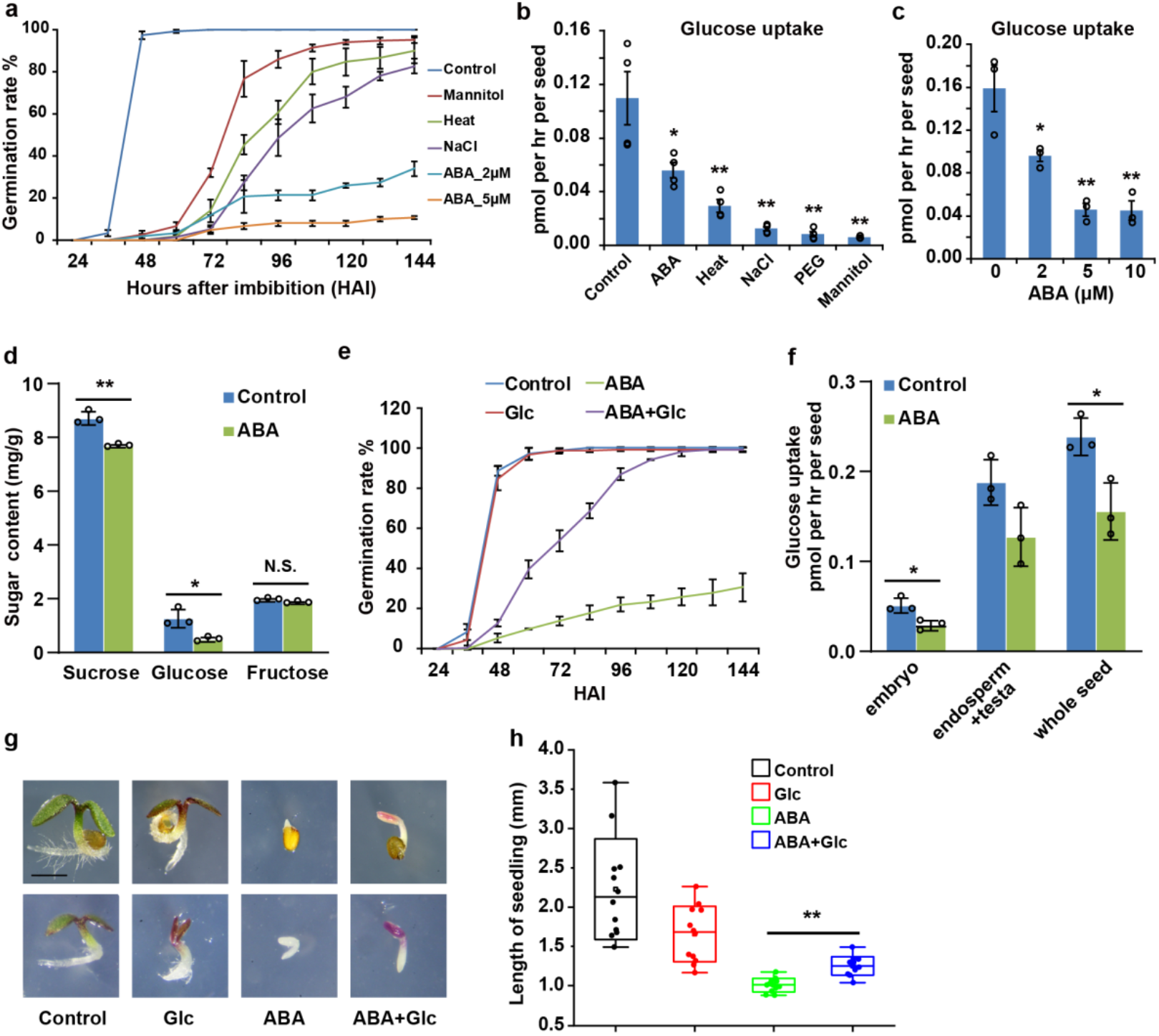
Abiotic stresses inhibit seed germination, while Glc relieves ABA inhibition. (*A*) Germination rate under different abiotic stress treatments. After 3 day imbibition, wild-type (Col-0) seeds were sown on ½ MS medium with 300 mM mannitol, 150 mM NaCl, and ABA (2 and 5 μM). Dry seeds pretreated at 50°C for 1 h before imbibition are shown as heat treatment. Values are means ± SEM (n = 3). (*B*) Glc uptake assay in seeds germinating under different abiotic stress treatments. Seeds were collected 24 hours after imbibition (HAI) and transferred to liquid ½ MS medium without (control) or supplemented with 2 μM ABA, 300 mM NaCl, 20% PEG_8000_, 600 mM mannitol, or incubated at 38 °C (heat) for 6 h. Values are means ± SEM (n = 4). (*C*) Glc uptake assay in seeds germinating under different concentrations of ABA. Seeds (24 HAI) were transferred to liquid ½ MS medium supplemented with 2, 5, or 10 μM ABA for 6-hour incubation. Values are means ± SEM (n = 3). (*D*) Sugar contents in ABA-treated seeds. After 3 day imbibition, wild-type seeds were incubated in liquid ½ MS medium without (control) or supplemented with 5 μM ABA for 24 hours. Values are means ± SD (n = 3). (*E*) Germination rate under Glc, ABA and ABA+Glc treatments. Values are means ± SD (n = 3). (*F*) Glc uptake in embryo, endosperm/testa and whole seed upon ABA treatment. The embryo and endosperm/testa were harvested separately after treatment with 2 μM ABA. Values are means ± SD (n = 3). (*G*) Representative images of 4-day-old WT seedlings with or without seed coat grown on medium supplemented with ABA, Glc or ABA+Glc. Scale bar, 1 mm. (*H*) Length of 4-day-old seedlings without seed coat as shown in (*G*). Embryos were dissected after imbibition and grown on the indicated medium. ABA suppresses embryo growth, while Glc can relieve this inhibition. Boxes are means ± SD with the median line (n = 12). The whiskers mark the minimum and maximum values. Asterisks indicate significant differences compared with control values (Student’s *t*-test, *p < 0.05; **p < 0.01). N.S., not significant.

As the endosperm provides nutrients and secretes hormones to transduce signals to the embryo and regulate embryo growth (Lee *et al*., 2010; Kang *et al*., 2015; Sanchez-Montesino *et al*., 2019), we measured Glc uptake in embryos and seed coats (endosperm + testa) separately. Interestingly, ABA exerted similar inhibition of Glc uptake into both embryo and endosperm (Fig. 1F), which indicates that Glc transporter(s) present on the outermost embryo cell layer and the single-cell layer of endosperm are both inhibited by ABA. The growth delay was similarly rescued by Glc in both dissected embryos and whole seeds (Fig. 1G), hinting that at least one Glc transporter on the plasma membrane of the embryo dermis moves the Glc into embryos to overcome ABA inhibition. These results also indicate that ABA inhibition of Glc import to embryos proportionally represents ABA inhibition of whole seeds (Fig. 1F and 1G).

### ABA restricts sugar availability in embryonic hypocotyl visualized by a FRET nanosensor

As hypocotyl elongation is indicative of high metabolic activity, we speculated that different tissues might differentially respond to ABA inhibition, resulting in the spatial distribution of soluble sugars, including Glc. To test this hypothesis, Glc FRET nanosensors were used to non-invasively monitor Glc levels (Fig. S3) at the subcellular level with high temporal resolution (Walia *et al*., 2018). We first determined which Glc sensor was suitable by comparing responses to the Glc available during seed germination using two cytosolic Glc sensors (in *rdr6*) substantially differing in their affinities (Deuschle *et al*., 2006). Due to challenges in imaging the embryo through the seed coat in an intact seed, we carefully isolated embryos at various time points until germinated, followed by microscopy imaging. As germination proceeded, cytosolic Glc levels increased, as indicated by darker colors from both the FLIPglu-170nΔ13 and the FLIPglu-3.2mΔ13 sensor (Fig. 2A) due to negative correlation between Glc level and the ratio of acceptor:donor emission for these two sensors. Since Glc levels progressively increase after imbibition (Pritchard *et al*., 2002), the high-affinity sensor FLIPglu-170nΔ13 was saturated about 12 hours before germination (Fig. 2A), while the FLIPglu-3.2mΔ13 was not saturated right through germination. Since we planned treatments with exogenous Glc in subsequent experiments, we decided to use the low-affinity sensor FLIPglu-3.2mΔ13 in Col-0 to monitor the spatiotemporal changes of Glc with different treatments during seed germination.

**Fig. 2.**
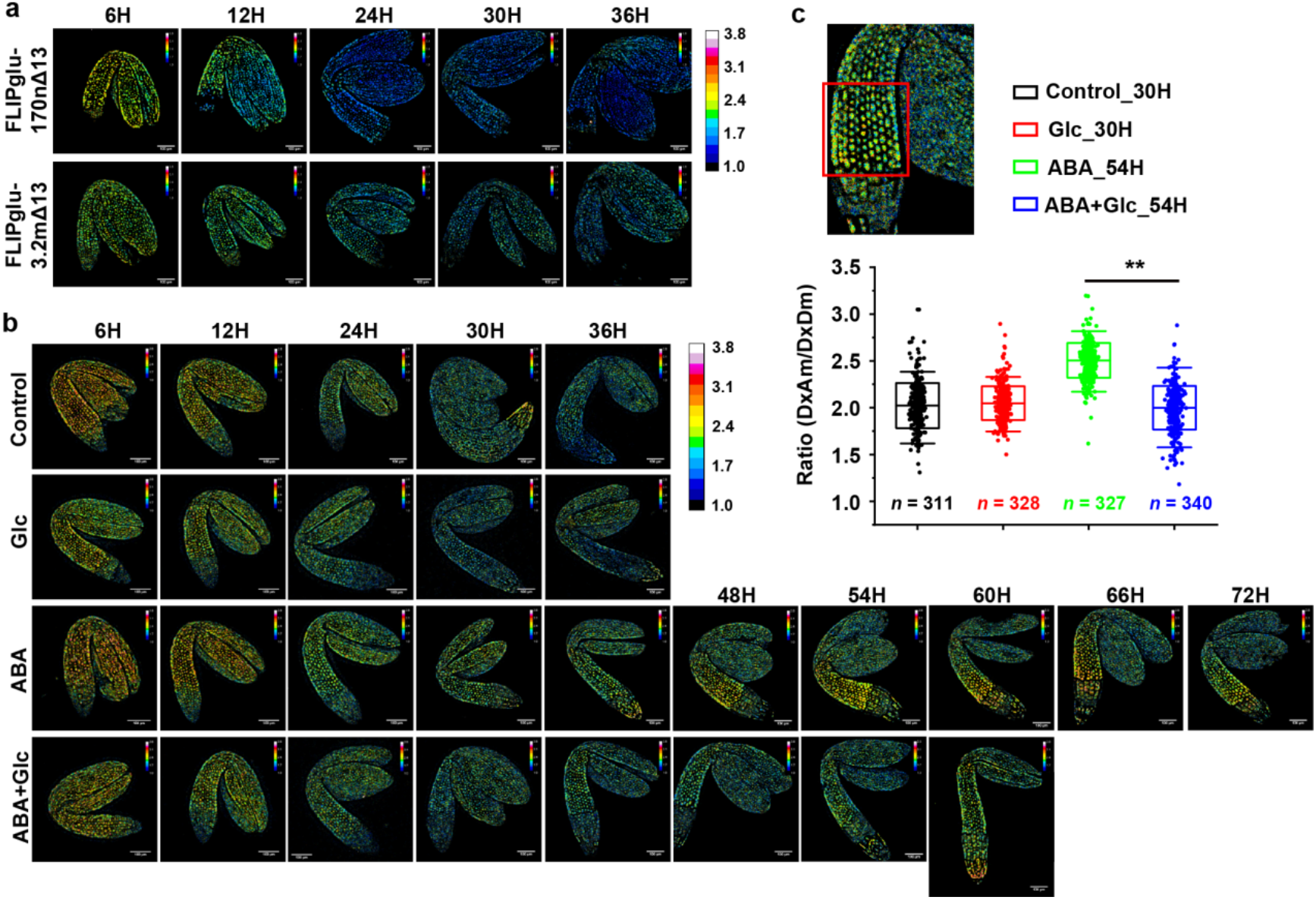
ABA restricts cytosolic Glc availability in the embryonic hypocotyl. (*A*) Glc nanosensors used in this study. Both sensors in *rdr6* show Glc levels increase after imbibition during seed germination. (*B*) FLIPglu-3.2mΔ13 sensor in Col-0 monitors the cytosolic soluble Glc level in embryo upon different treatments of germinating seeds. The ratio of acceptor emission (Am) to donor emission (Dm) under donor excitation (Dx) is shown, with each row representing a different treatment, at the indicated time points until seeds germinated (the last embryo of each row). The DxAm/DxDm ratio negatively correlates with Glc concentration. ABA restricts Glc availability in lower hypocotyl compared to other treatments, while exogenous Glc can partially rescue this restriction, as shown in the bottom row. Scale bar = 100 μM. (*C*) Quantification of DxAm/DxDm ratio before germination shown in (*B*) in the embryonic hypocotyl region upon treatments. The region of interest is marked with a red frame in the upper image. Boxes represent mean ± SD with the median line (n > 300). The whiskers mark the outliers. Asterisks indicate significant differences (Student’s *t*-test, **p < 0.01).

We compared dynamic Glc distribution in embryos dissected from ABA-treated FLIPglu-3.2mΔ13 seeds with that from non-treated seeds. In ABA-treated seeds, the overall progression of Glc accumulation was slowed (Fig. 2B). Before the transition from seed to seedling, Glc level was dramatically reduced in the hypocotyl region with ABA treatment, as indicated by the brighter colors (Fig. 2A), and the diminished Glc levels extended from the hypocotyl to the radicle until the radicle emerged. Excitingly, higher Glc was detected in the hypocotyl region of seeds treated with both ABA and Glc (ABA+Glc) than in seeds treated with ABA only, with the most remarkable difference at the transition stage from seed to seedling, as quantified in (Fig. 2C). This is consistent with the observation that Glc can override ABA-seed inhibition to a certain degree (Fig. 1E).

### Sugar metabolism associated genes are highly enriched under ABA treatment

At any given time, a FRET sensor detects the steady state level of a ligand, which results from influx, efflux and metabolism. This particular cytosol-localized Glc sensor monitors the cytoplasmic Glc pool resulting from Glc exchange across the plasma membrane, from subcellular compartmentation of Glc mediated by the sugar transporters, and the combined effects of enzyme-directed sugar metabolism. Our quantification data of Glc distribution upon ABA treatment showed that ABA controls overall and regional availability of Glc, providing energy for many biological events during seed germination (Al-Ani *et al*., 1985). To explore the possible mechanism responsible for ABA regulation of cytosolic Glc, transcriptome analysis was used to mine for genes responding to Glc, ABA and ABA+Glc, and to find possible regional players by combining gene expression profiles from different seed compartments (Dekkers *et al*., 2013).

WT seeds at 24 hours after imbibition (HAI) were treated with Glc, ABA, or ABA+Glc for 6 hours and the total RNA was extracted from whole seeds for RNAseq. Through pairwise comparison, 5112 differentially expressed genes (DEGs, |b| ≥ 0.2, q < 0.05) were identified among the treatments. Genes that primarily function in pathways such as glycerolipid metabolism, starch and sucrose metabolism, and glycolysis/gluconeogenesis were highly enriched (Table S1), consistent with our findings that ABA changes sugar levels in germinating seeds (Fig. 1D and Fig. 2B). The heatmap shows that the DEGs were grouped into four clusters based on similar expression patterns (Fig. 3A and Table S2). Since we aimed to explore the mechanism(s) governing inhibition of Glc distribution by ABA and antagonism of the inhibition by Glc, we primarily focused on those genes differentially expressed among control, ABA and ABA+Glc. Gene ontology (GO) analysis showed that genes induced by ABA and to a lesser degree by ABA+Glc in cluster 1 are mostly involved in water deprivation response, alcohol metabolism, seed dormancy, fatty acid catabolism, lipid storage, or responses to other various stresses, and in cluster 4 function in biological processes mainly associated with amine catabolism, karrikin signals, cell wall organization and biogenesis, and intracellular signal transduction.

**Fig. 3.**
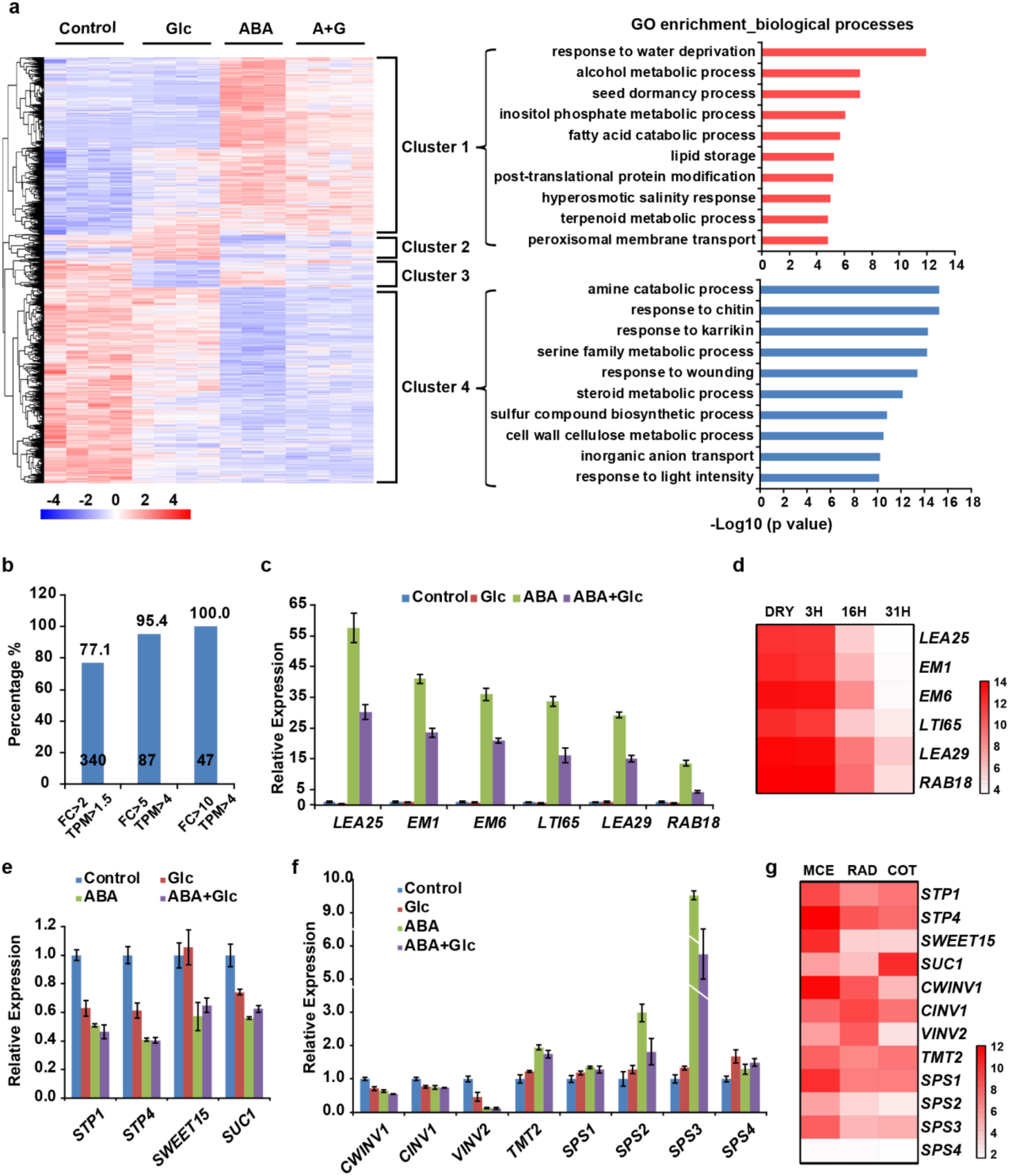
ABA suppresses sugar allocation and metabolism. (*A*) Heatmap of differentially expressed genes (DEGs). Four clusters of DEGs have been classified based on the co-expression pattern. The 10 most significantly enriched GO terms in cluster 1 and 4 are presented with brackets. (*B*) Glc compromises ABA response globally. The percentage of ABA-induced genes weakened by Glc correlates to the ABA induction level. (*C*) ABA marker genes are highly induced by ABA, and their expression was reduced by addition of Glc. (*D*) Heatmap showing levels of ABA marker genes shown in (*C*) in whole dry seeds (DRY) and germinating seeds at 3, 16 and 31 HAI (Dekkers *et al*., 2013). The levels of these genes declined as seed germination progressed. (*E*) ABA suppresses expression of sugar transporters. *STP1*, *STP4* and *SUC1* are suppressed by both Glc and ABA. *SWEET15* is not suppressed by Glc. (*F*) Sugar metabolism-related genes are regulated by ABA. Invertase genes *CWINV1*, *CINV1* and *VINV2* are down-regulated by ABA. *TMT2* and all four *SPS* genes are activated by ABA. (*G*) Expression of the genes shown in (*F*) in micropylar and chalazal endosperm (MCE), radicle (RAD) and cotyledons (COT) at 31 HAI. The RAD includes hypocotyl region as described (Dekkers *et al*., 2013).

### ABA response is compromised globally by addition of Glc

There is a total of 3156 genes were differentially expressed upon ABA+Glc treatment compared to control condition, 44% of which overlapped with genes regulated by ABA only. Interestingly, among the 340 ABA-induced genes (FC > 2, TPM > 1.5), 262 genes (77.1%) were reduced by ABA+Glc compared to ABA (Fig. 3B). Among the ABA-induced genes (with induction greater than 5- and 10-fold), 95.4% and 100% were lessened by addition of Glc, respectively (Fig. 3B), suggesting that Glc compromises the ABA response globally. *LEA25*, *LEA29*, *EM1*, *EM6*, *LTI65* and *RAB18*, as ABA-responsive marker genes, were dramatically induced by ABA, but only reached 40-60% of these levels with addition of Glc (Fig. 3C), indicating that down-regulation of their expression is associated with the completion of germination. Consistent with this observation, the expression of the top 50 ABA-inducible genes (TPM > 1.5), including these six marker genes mentioned above, declined over the course of germination under normal conditions (Fig. 3D and Fig. S4). The addition of Glc likely relieves the suppression imposed by ABA through down-regulating ABA-responsive genes, including dormancy-associated genes (Bassel *et al*., 2011) (Table S3). The reduced uptake and cytosolic accumulation of Glc in ABA-treated seeds are a cumulative result of (1) impaired Glc influx across the plasma membrane, as ABA suppressed sugar transporter genes such as endosperm enriched *STP1*, *STP4*, *SWEET15* and cotyledon enriched *SUC1* (Fig. 3E and 3G); (2) enhanced Glc import into the vacuole, as ABA up-regulated the expression of *TMT2*, a tonoplast localized proton/monosaccharide antiporter; and (3) decreased production of Glc, as ABA down-regulated sucrose cleavage by invertases such as *CWINV1*, *CINV1* and *VINV2* highly accumulating in hypocotyl and radicle (Fig. 3F and 3G). It is interesting that the *SPS* genes were up-regulated by ABA (Fig. 3F), which indicates that more hexose is converted to sucrose, resulting in lower Glc levels.

### The *sps* mutants accumulate more Glc in seeds and germinate earlier under ABA treatment

SPS is a rate-limiting enzyme for sucrose synthesis using UDP-glucose and fructose 6-phosphate. In *sps* null mutants, plants accumulate more Glc in leaves (Volkert *et al*., 2014; Bahaji *et al*., 2015). To genetically evaluate if *SPS* genes contribute to Glc availability in germinating seeds, we measured the sugar contents in *sps* mutant seeds, and found no significant difference in sucrose levels in *sps1*,*4* mutant, less sucrose in *sps1*,*2*,*3* mutant (Fig. 4A), but higher Glc and fructose in both *sps* mutants (Fig. 4B). As expected, the *sps* seeds germinated earlier under ABA treatment, and *sps* seedlings were less sensitive to ABA (Fig. 4C). Interestingly, not only were the *SPS* expressions altered, but so were many other enzymes involved in starch and sucrose metabolism (Fig. S5). These data indicate that higher endogenous sugar levels facilitate earlier seed germination under ABA, and likely also under other stressed conditions.

**Fig. 4.**
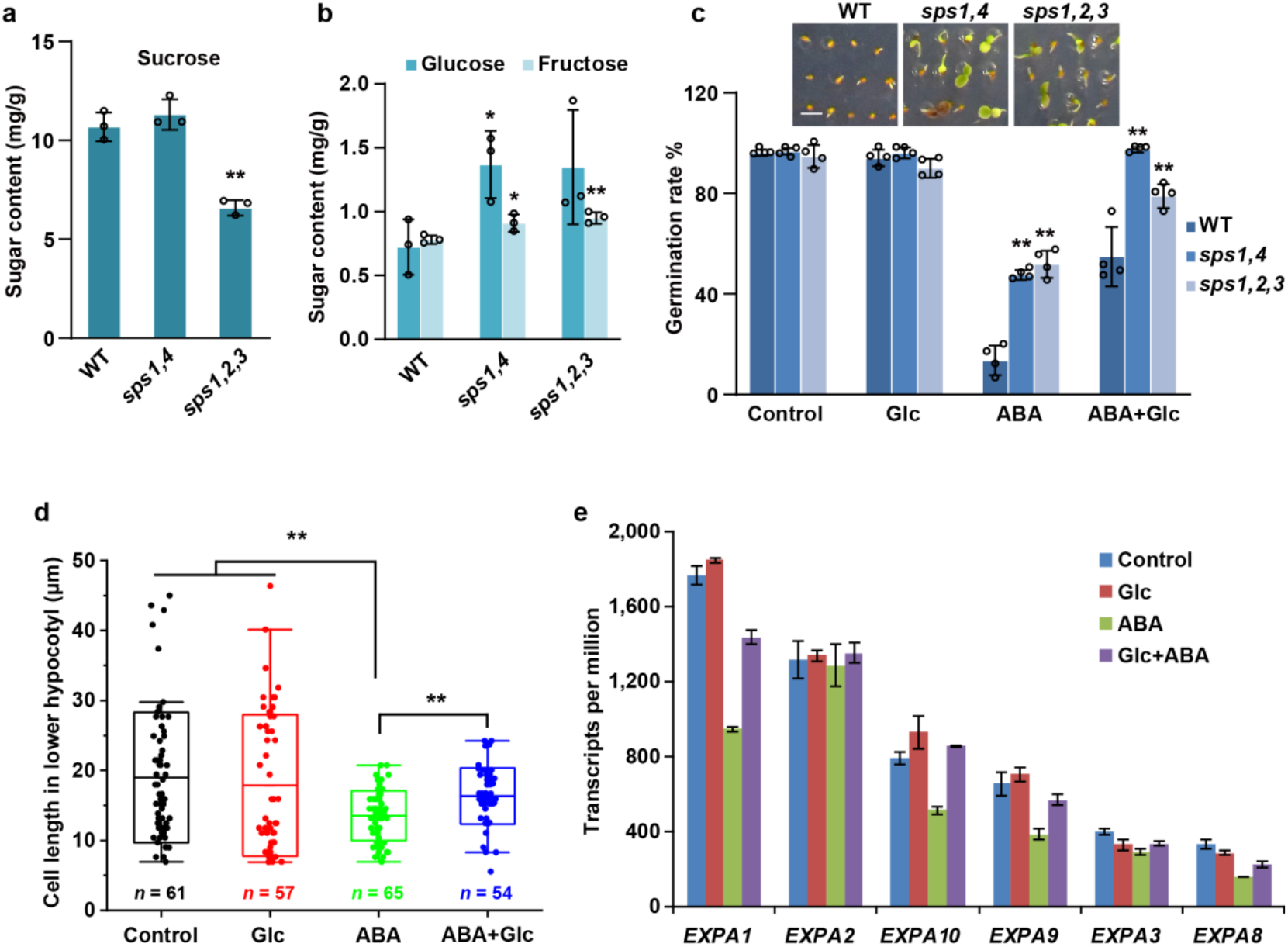
Glc relieves ABA-triggered germination delay through reactivating *EXPA* gene family. (*A* and *B*) Sugar contents in *sps* mutant seeds during germination. Sugar contents are measured at 30 HAI. There is less sucrose in *sps1,2,3* (*A*), but more Glc and fructose in both *sps1,4* and *sps1,2,3* seeds (*B*). Values are means ± SD (n = 3). (*C*) The *sps* mutants are less sensitive to ABA. Germination rates of *sps* mutants under Glc, ABA and ABA+Glc treatments were calculated at 96 HAI. Values are means ± SD (n = 4). The inset images on top show the representative 14-day-old seedlings grown on medium supplemented with 2 μM ABA. Scale bar, 2 mm. (*D*) Length of epidermal cells in embryonic hypocotyl upon treatment. Cell length of lower hypocotyl region adjacent to transition zone was measured upon Glc, ABA and ABA+Glc treatment at 36 HAI. Boxes represent mean ± SD with the median line (n > 50). The whiskers mark the outliers. (*E*) ABA suppresses *EXPA* expression. The six most abundant *EXP* genes are suppressed by ABA, except *EXPA2*. Glc counters the ABA suppression. Asterisks indicate significant differences compared with corresponding WT values (Student’s *t*-test, *p < 0.05; **p < 0.01).

### Glc reactivates the elongation of hypocotyl cells by releasing the inhibition of ABA on *EXP* family

The completion of seed germination is attributed to hypocotyl elongation (Sliwinska *et al*., 2009). Genes related to cell wall synthesis or remodeling, such as glycosidases, peptidases and EXPs, are critical for germination (Mikola & Kolehmainen, 1972; Leah *et al*., 1995; Bassel *et al*., 2014; Xu *et al*., 2020). The genes encoding glycosidases, such as *XTH5* and *XTH31*, as well as peptidases, like *SCPL20*, *SCPL44* and *SCPL48*, were suppressed by ABA but activated by addition of Glc (Fig. S6). KEGG analysis found that cluster 4 was highly represented by pathways associated with amino sugar and nucleotide sugar metabolism, interconversions that regulate carbon flux into plant cell walls (Seifert, 2004; Bar-Peled & O’Neill, 2011) (Fig. S7). Measurements of cell length in the lower hypocotyl region revealed that Glc aided the cell elongation under ABA application (Fig. 4D), an observation that coincided with ABA-based suppression of the six most abundant *EXP* genes in germinating seeds, which was relieved by addition of Glc for all except *EXPA2* (Fig. 4E and Fig. S8). Thus, ABA likely limits essential nucleotide-sugar metabolism and/or partitioning to hamper cell wall expansion, thereby suppressing seed germination, as recently shown that EXPA-mediated cell wall loosening promotes germination under GA deficiency (Xu *et al*., 2020). Remarkably, all these findings align with the conclusion that Glc relieves the ABA inhibition on cell elongation in the lower hypocotyl through countering ABA suppression of downstream *EXPA* genes.

## Discussion

### ABA delaying seed germination is a result of inhibiting multiple pathways and Glc relieves the inhibition partially

Glc can, but not fully relieve ABA inhibitory effects on germination (Fig. 1E), which is consistent with the partial rescue of Glc spatiotemporal distribution by Glc in the ABA-treated versus untreated seeds as detected by a FRET Glc sensor (Fig. 2B). A partial explanation is that and Glc cannot fully overcome ABA-regulated gene expression especially for sugar transporters and sugar metabolism associated enzymes, such as STP members and invertases (Fig. 3E and 3F). These results indicate inhibition of germination resulting from ABA treatment is not simply due to limited Glc availability. Peptone has been reported to relieve ABA inhibition of germination (Garciarrubio *et al*., 1997), indicating that deficiency of amino acids may be triggered by ABA resulting in germination delay. In addition to nutrient deficiency, other hormone signaling pathways, such as gibberellin and ethylene, are suppressed by ABA (Ghassemian *et al*., 2000; Seo *et al*., 2006). Therefore, gibberellin and ethylene can antagonize ABA effects during germination (Beaudoin *et al*., 2000; Lee *et al*., 2010). The interaction between sugars and other hormones has been reported (Leon & Sheen, 2003; Matsoukas, 2014). It is worth noting that exogenous Glc can impair the functions of ethylene (Yanagisawa *et al*., 2003; Price *et al*., 2004) and gibberellin (Perata *et al*., 1997; Yuan & Wysocka-Diller, 2006). So Glc cannot completely rescue the germination delay caused by ABA may be due to ABA-antagonizing hormone signaling, such as gibberellin and ethylene, are repressed by Glc itself. However, we cannot exclude the possibility that some modifications to transporters, enzymes or signaling components at the post-translational level imposed by ABA are unable to be reversed by the addition of Glc, although no changes in expression are observed.

### Glc and Fru are more effective than Suc to antagonize ABA during seed germination

Metabolizable sugars (Suc, Glc and Fru) can relieve the ABA inhibition on germination (Garciarrubio *et al*., 1997; Finkelstein & Lynch, 2000). But exogenous Fru is less effective than same concentration of Suc and Glc (Finkelstein & Lynch, 2000), which may be due to the lack of an efficient Fru transporter in germinating seeds. The *sps1,4* mutant with unchanged Suc and *sps1,2,3* with less Suc, both containing higher levels of Glc and Fru, germinated earlier than WT upon ABA treatment (Fig. 4C). These data suggested that Glc and Fru, but not Suc, are sufficiently antagonistic to ABA *in vivo* during germination, and exogenous Suc may require the efficient break down into Glc and Fru by invertases or sucrose synthase for sufficient antagonistic role to ABA, implied by the finding that the same concentration (w/v) of exogenous Glc is more effective than Suc in antagonizing 10 μM ABA (Garciarrubio *et al*., 1997).

### Hypocotyl is the most sensitive subdomain in the embryos in response to environmental stimuli during seed germination

The lower hypocotyl and hypocotyl–radicle transition zone has been identified as the region responsible for cell elongation to complete germination in Arabidopsis (Sliwinska *et al*., 2009). Our FRET sensor imaging data coincidentally revealed that ABA inhibited the Glc accumulation in the lower hypocotyl, but not in the cotyledon or radicle (Fig. 2B). These data are consistent with the previous findings that the hypocotyl is the site to be induced by red light during germination (Inoue & Nagashima, 1991). Interestingly, complex sugars and hormones also highly accumulate in the embryonic hypocotyl before germination. For example, many carbohydrate-containing granules (starch or other glucans) appear in the lower hypocotyl region just before germination (Sliwinska *et al*., 2009) and the key gibberellin biosynthesis genes *GA3ox1* and *GA3ox2* are specifically active in the hypocotyl (Yamaguchi *et al*., 2001). In addition, Stamm *et al*. quantified the single-cell 3D growth over germination and found that gibberellin-ATHB5-EXP3 cascade mediated longitudinal cell growth in the hypocotyl is responsible for the completion of germination (Stamm *et al*., 2017). Therefore, light, gibberellin, sugar and ABA converge on hypocotyls to control seed germination. Our results further proved that the hypocotyl is the most important subdomain in the embryos in response to environmental stimuli during seed germination. The single-cell sequencing technology may be used to explain why hypocotyl is the most active tissue to control seed germination. In combination with other hormone bio-nano sensors with high resolution, such as the GA sensor (Rizza *et al*., 2017) or the ATP sensor (De Col *et al*., 2017), can help to better understand the molecular control in the hypocotyl for seed germination.

### Glc and ABA antagonize each other in multiple dimensions

Glc serves not only as a nutrient molecule, but as a signal molecule mediated either by Hexokinase 1 (HXK1) (Jang *et al*., 1997; Moore *et al*., 2003), the regulator of G-protein signaling 1 (RGS1) (Chen *et al*., 2003), or target of rapamycin (TOR) (Menand *et al*., 2002; Xiong & Sheen, 2012). Wang *et al*. found that Glc-activated TOR kinase phosphorylates ABA receptors to prevent activation of ABA signaling under optimal conditions, whereas ABA-activated SnRK2s phosphorylate Raptor, a component of TOR complex in plants, to inhibit TOR signaling under stress conditions (Wang *et al*., 2018). But it is not clear that if TOR signaling is activated under ABA+Glc treatment during germination. The ABA-mediated restriction of sugar in the hypocotyl partially explains ABA-delayed seed germination, as discussed above. This restriction is mainly achieved by regulating the expression of sugar transporters and sugar metabolism-related enzymes (Fig. 3E and 3F). Notably, we found that several ABA receptors were down-regulated by exogenous Glc during germination (Fig. S9). Therefore, Glc and ABA antagonize each other at both transcriptional and post-translational levels. However, we cannot exclude the possibility that Glc directly inactivates the ABA, such as reduces the active form of ABA via ABA-glucose ester formation (Boyer & Zeevaart, 1982), which deserves further investigation.

## Acknowledgments

We thank the former member of the Wolf Frommer lab, Bi-Huei Hou, for generating the Ubi10-FLIPglu-3.2mΔ13 sensor in Col-0. We thank Wolf Frommer at Heinrich Heine University Düsseldorf for sharing glucose sensor seeds used in this research. We appreciate Javier Pozueta-Romeroa at Agrobiotechnology Institute (IdAB) for providing the *sps* mutant seeds. We thank Ching Man Wai for the initial analysis of RNA-seq data. This work was supported by start-up funds from the University of Illinois at Urbana-Champaign to Dr. Li-Qing Chen.

## Author contributions

XX, Y-CY, and L-QC planned and designed the experiments. XX and Y-CY conducted the experiments and XX, Y-CY, YW, HX and L-QC analyzed the data. XX, Y-CY, and L-QC wrote the manuscript.

## Data availability

All the data and materials that support the findings of this study are available upon request from the corresponding author. The RNA-sequencing data described in this study has been deposited to the National Center for Biotechnology Information GEO database (accession number GSE163057).

## Supplementary Figures

**Fig. S1.**
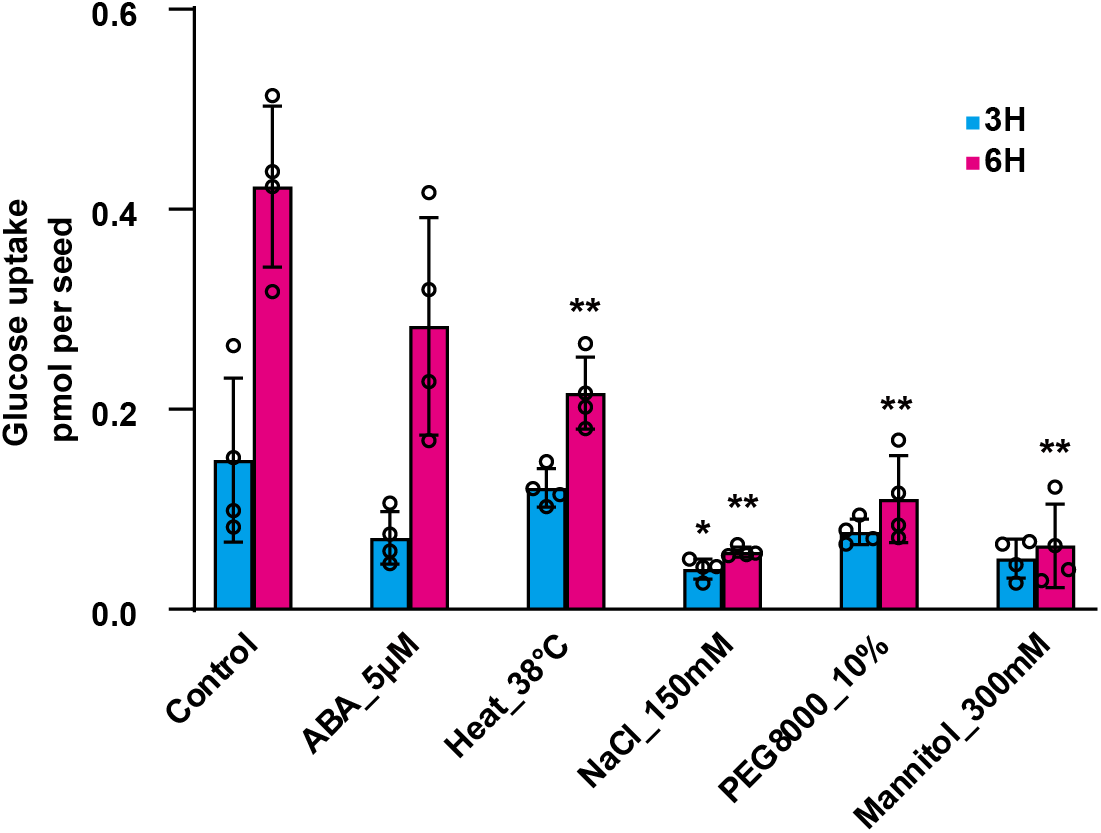
Abiotic stresses suppress Glc sugar uptake during seed germination. Glc uptake was suppressed by abiotic stresses in germinating seeds. Seeds (24 HAI) were collected and transferred to liquid ½ MS medium without (control) or supplemented with 2 μM ABA, 150 mM NaCl, 10% PEG_8000_, or 300 mM mannitol and incubated at 22 °C or kept in control media and incubated at 38 °C (heat) for 3 h and 6 h. Values are means ± SD (n = 4). Asterisks indicate significant differences compared with control values (Student’s *t*-test, *p < 0.05; **p < 0.01).

**Fig. S2.**
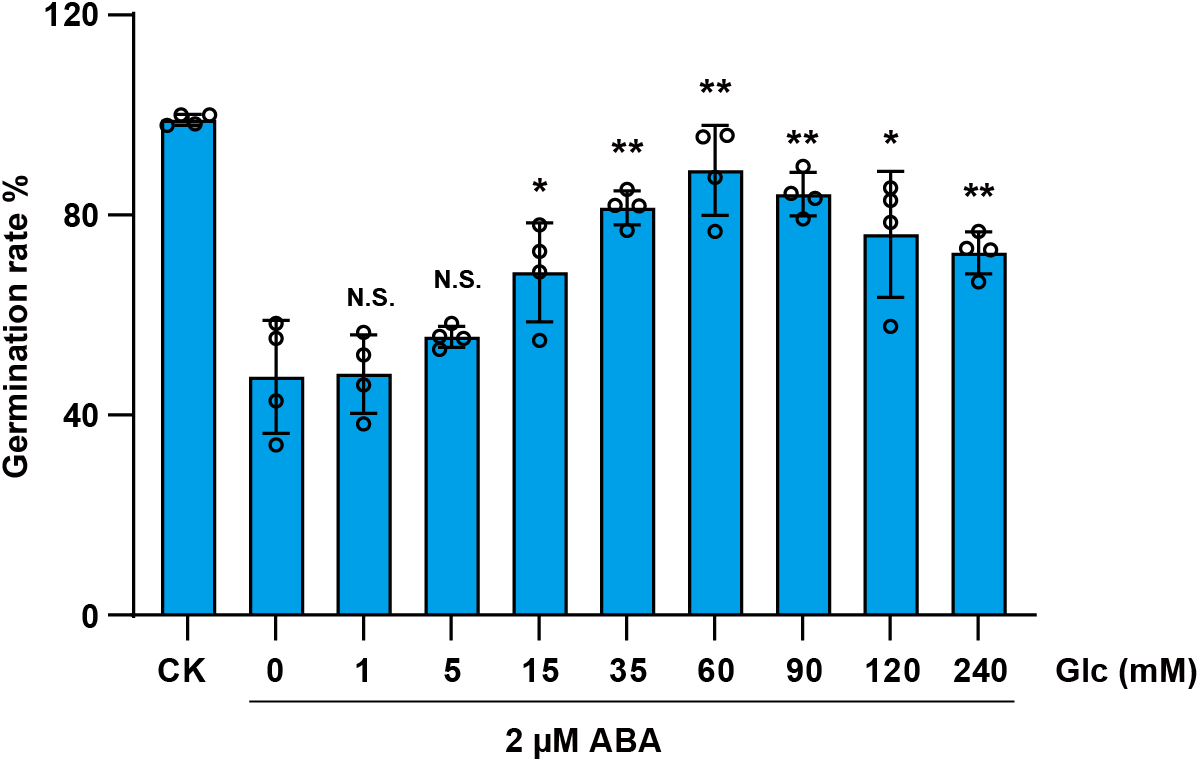
In a concentration-dependent manner Glc relieves ABA inhibition on germination. In the presence of ABA, the germination rate dropped without Glc, but nearly recovered at 60 mM Glc, although 15 mM Glc was effective. WT seeds were grown on medium supplemented with 2 μM ABA and the germination rate was calculated at 72 HAI. Values are means ± SD (n = 4). Asterisks indicate significant differences compared with ABA values (Student’s *t*-test, *p < 0.05; **p < 0.01). N.S., not significant.

**Fig. S3.**
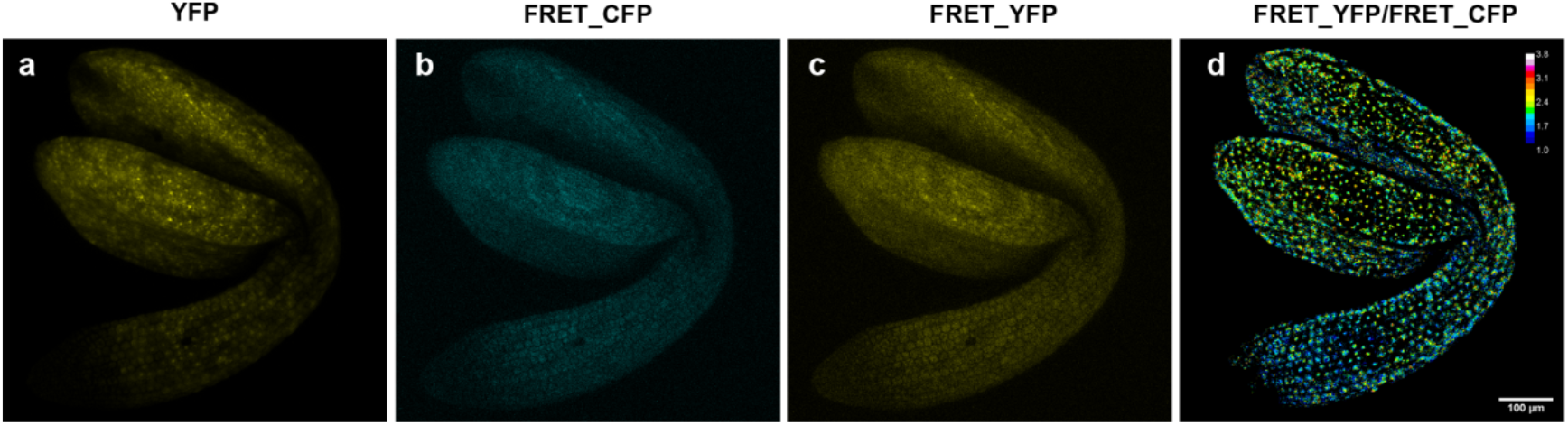
Imaging embryo with FLIPglu-3.2mΔ13 Glc sensor. (*A*) Image was taken under acceptor (YFP) excitation, 514 nm. (*B*) FRET_CFP image was taken under donor (CFP) excitation, 458 nm. (*C*) FRET_YFP image was taken under donor (CFP) excitation, 458 nm. (*D*) An image of FRET ration. FRET_CFP and FRET_YFP were used to calculate the ratio, which was processed and analyzed as described (Jones *et al*., 2014).

**Fig. S4.**
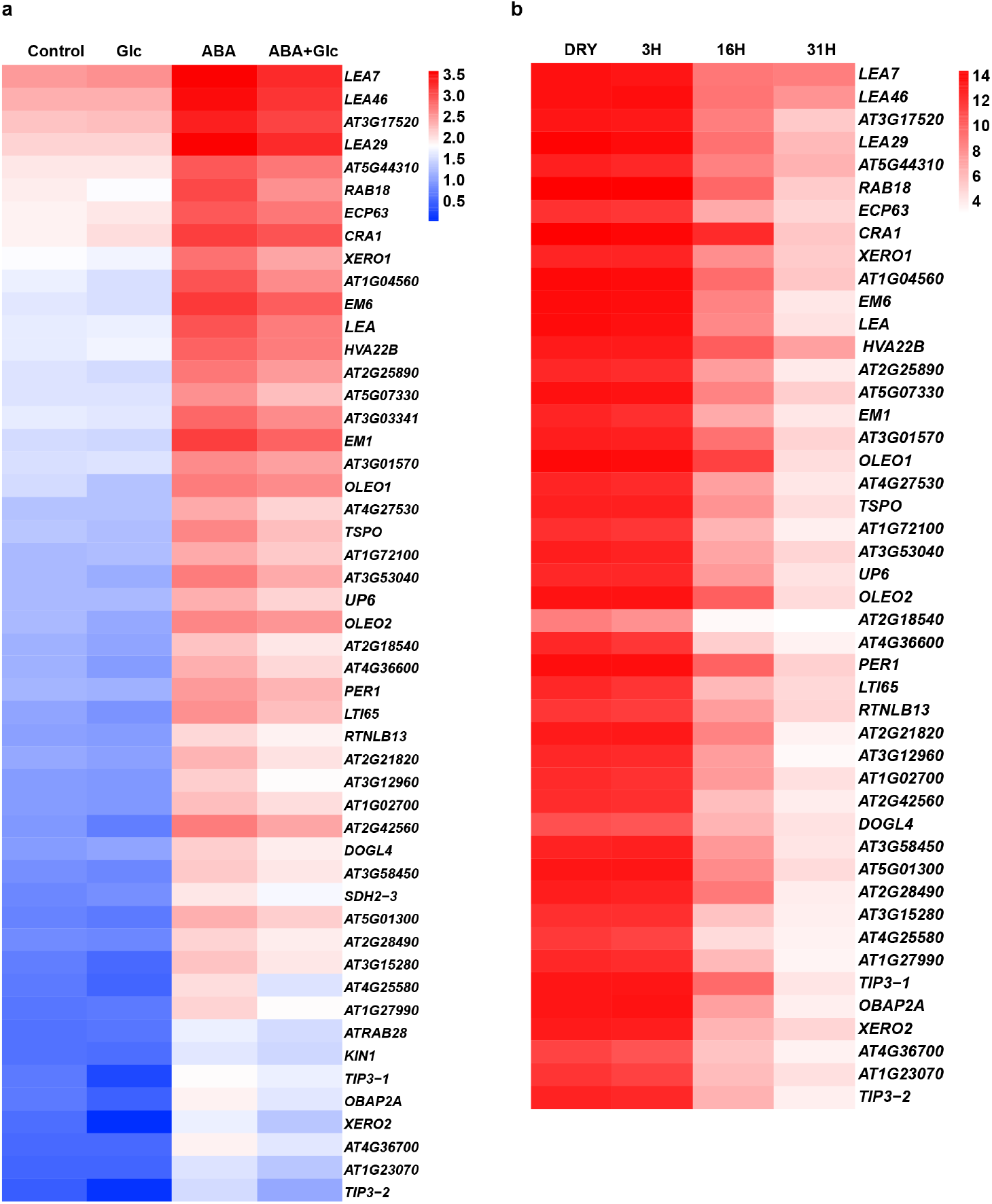
Glc compromises dormancy-related gene expression. (*A*) Heatmap shows the expression of the top 50 ABA-inducible genes with TPM>1.5 under our treatment conditions. The degree of induction is weakened by ABA+Glc compared to ABA. (*B*) Heatmap shows levels of ABA-inducible genes shown in (*A*) in whole dry seeds (DRY) and germinating seeds at 3, 16 and 31 HAI (Dekkers *et al*., 2013). The levels of 46 genes are available in the dataset (Dekkers *et al*., 2013). The average values of the three tissues (endosperm, radicle and cotyledon) at each time point are calculated to make this heatmap. The levels of all these genes declined as seed germination progressed.

**Fig. S5.**
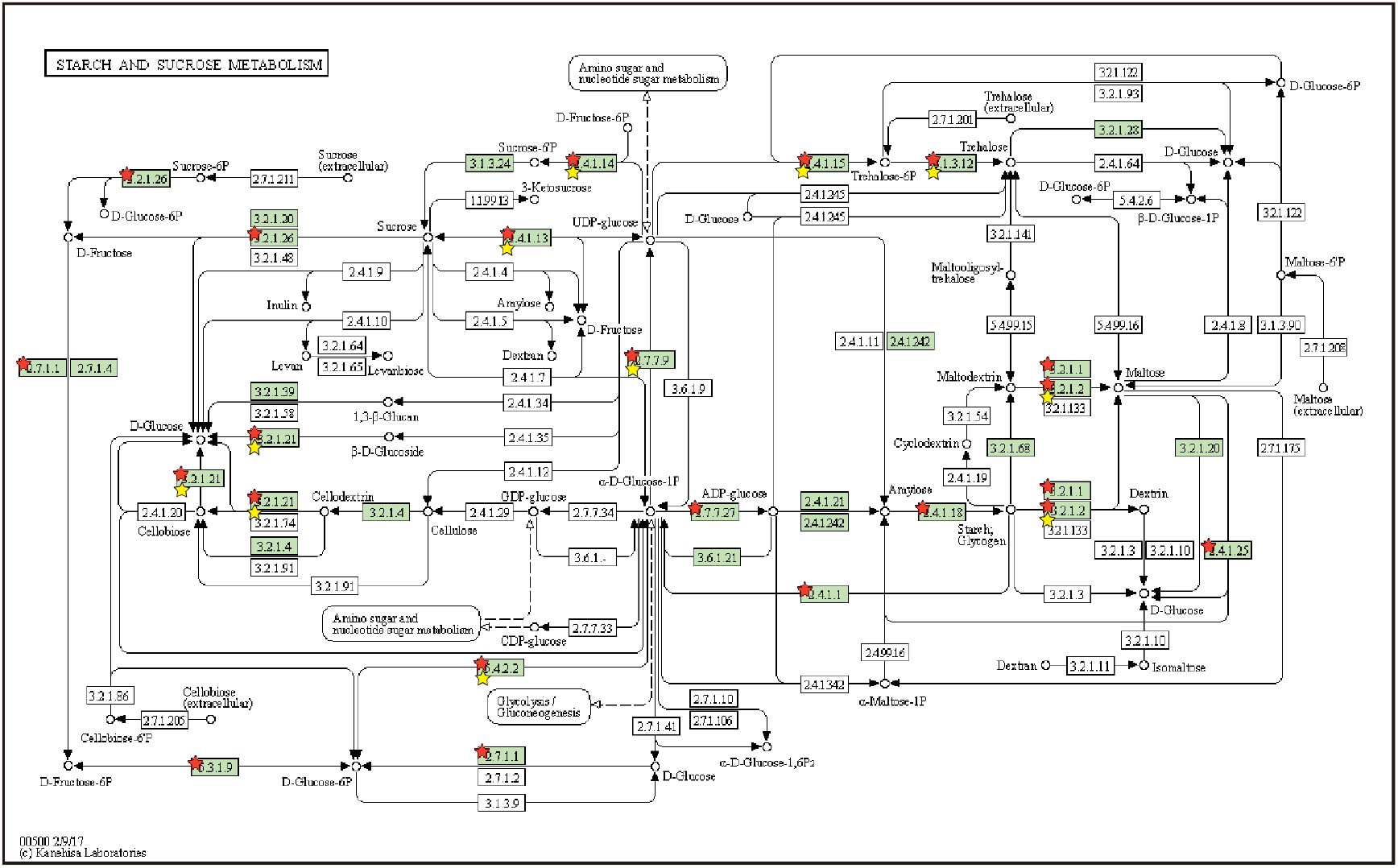
Genes in the starch and sucrose metabolism pathways are up-regulated by ABA. KEGG pathway was analyzed using DAVID Bioinformatics Resources 6.8 (Huang da *et al*., 2009). Genes up-regulated by ABA are labeled with red stars, while the genes suppressed by ABA+Glc compared to ABA are marked with yellow stars.

**Fig. S6.**
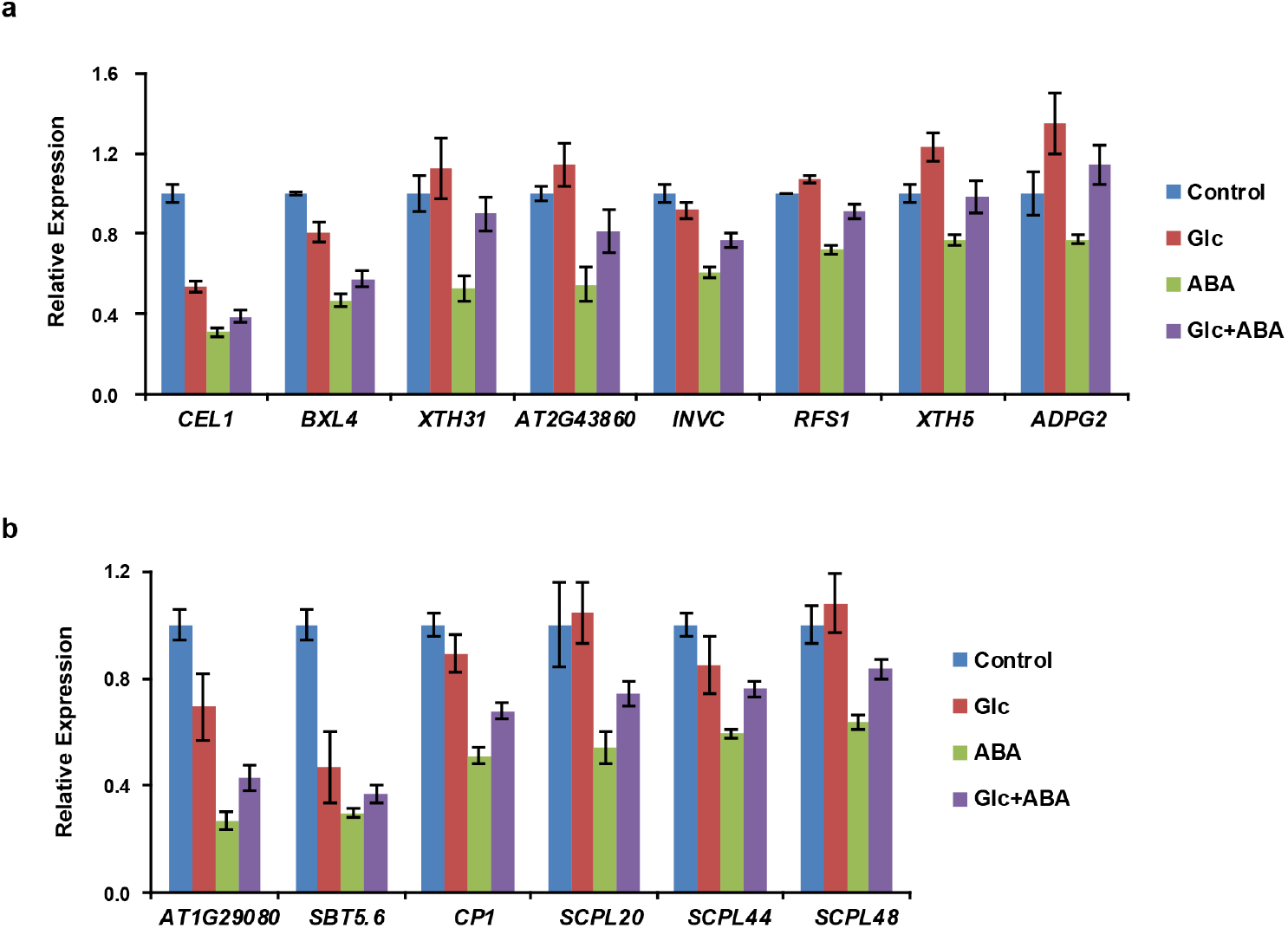
ABA suppressed genes encoding glycosidases and peptidases, but Glc relieved this suppression. The glycosidases hydrolyze complex sugars into small carbon skeletons, while the peptidases cleave proteins into smaller polypeptides or amino acids, providing substrates for the re-synthesis of oligo- and poly-saccharides. (*A*) Glycosidase encoding genes *CEL1*, *BXL4*, *XTH31*, *AT2G43860*, *INVC*, *RFS1*, *XTH5* and *ADPG2* were suppressed by ABA, while Glc addition can relieve this suppression such that some can reach the levels without treatment, such as *XTH5* and *ADPG2*. (*B*) Peptidase encoding genes *AT1G29080*, *SBT5.6*, *CP1*, *SCPL20*, *44* and *48* were suppressed by ABA.

**Fig. S7.**
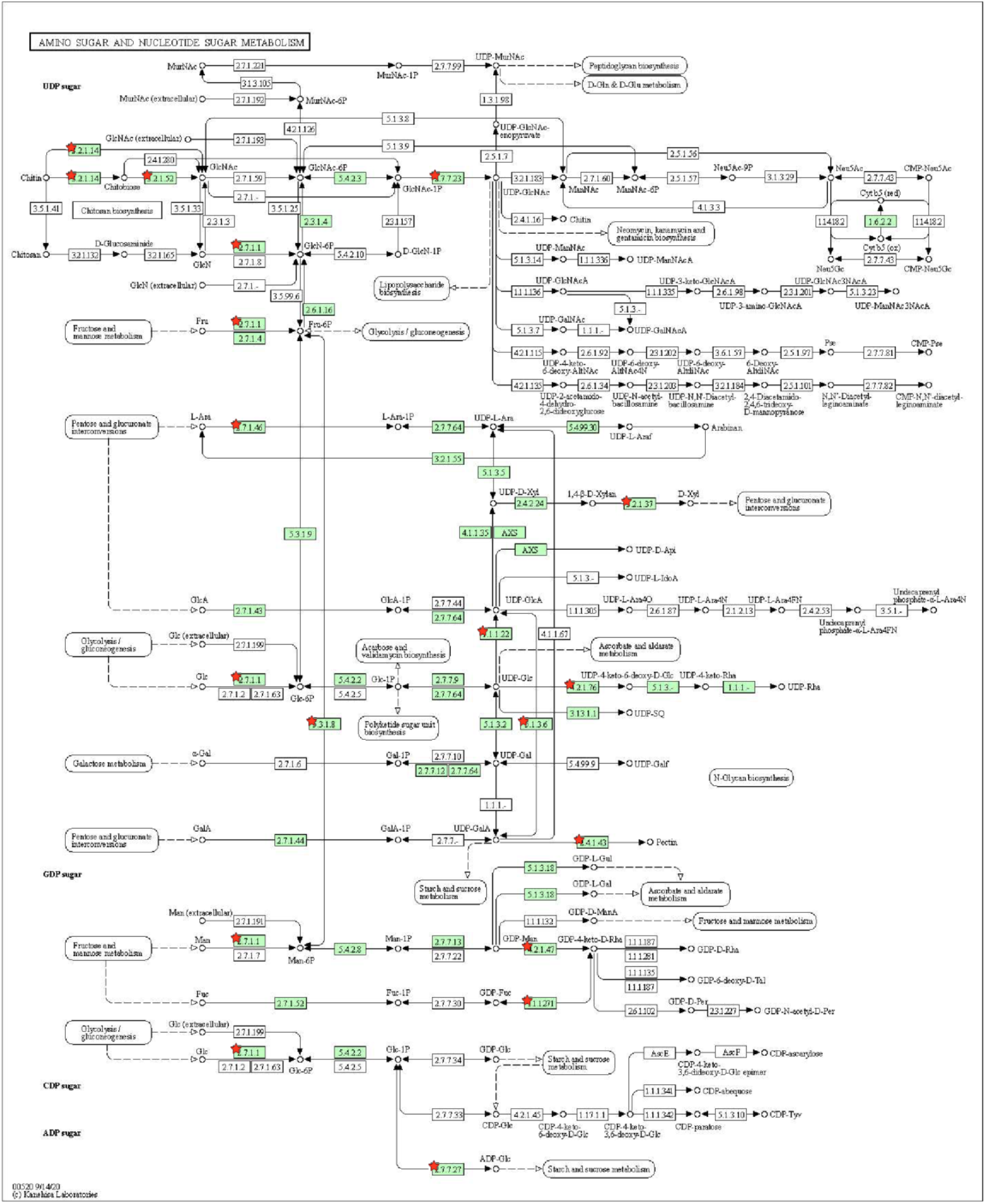
Genes in the amino sugar and nucleotide sugar metabolism pathways are down-regulated by ABA. KEGG pathway was analyzed using DAVID Bioinformatics Resources 6.8 (Huang da *et al*., 2009). Genes suppressed by ABA are labeled with red stars.

**Fig. S8.**
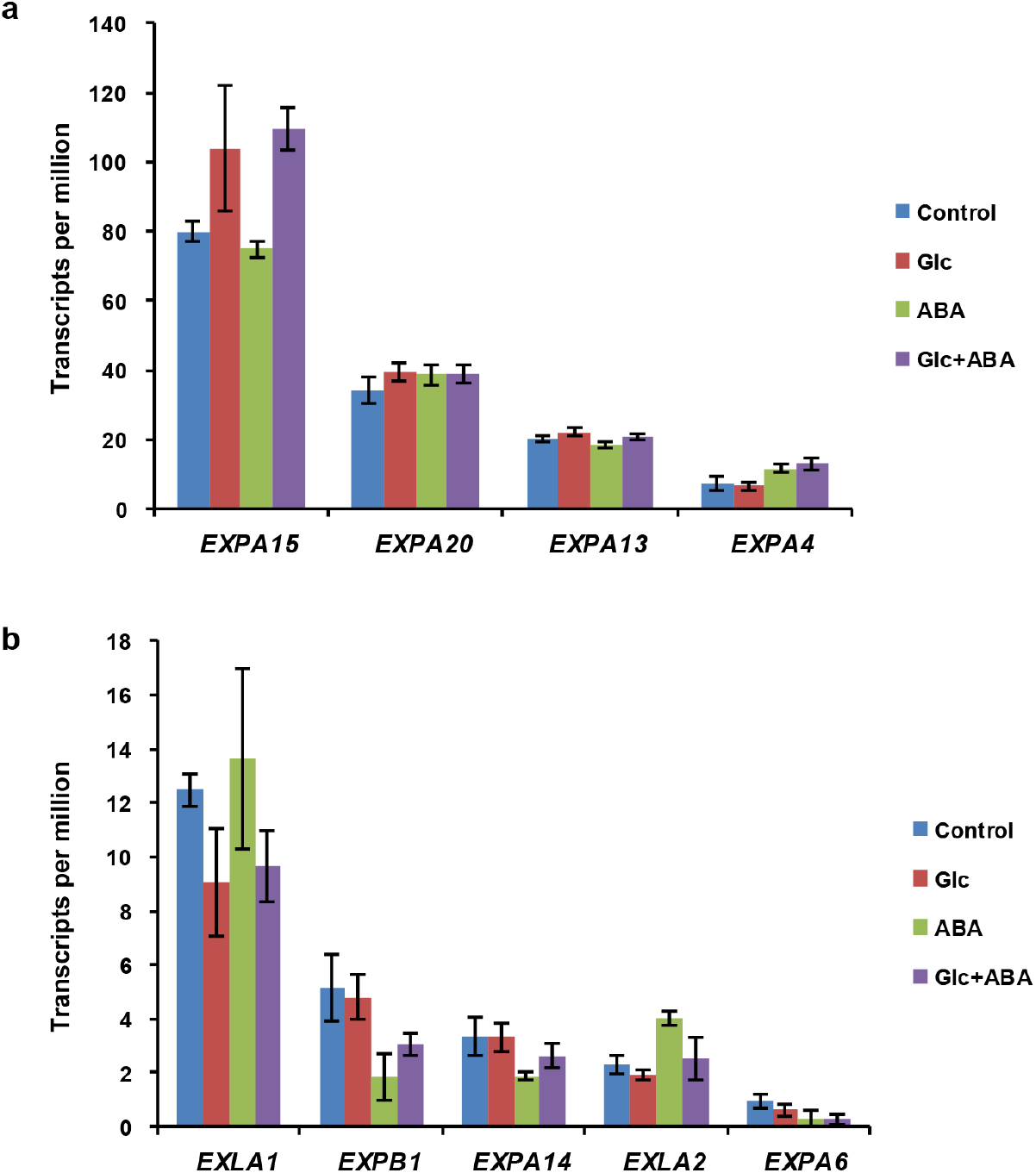
Expression of low-abundance *Expansin* genes under treatments. 15 *Expansin* (*EXP*) genes are active during seed germination (TPM > 1). The levels of most six abundant genes (TPM > 300) were shown in Fig. 4E. (*A*) Expression of genes *EXPA15*, *20*, *13* and *4* (15 < TPM< 80) were not suppressed by ABA. (*B*) Levels of genes *EXLA1*, *EXPB1*, *EXPA14*, *EXLA2* and *EXPA6* (TPM < 15) under treatment.

**Fig. S9.**
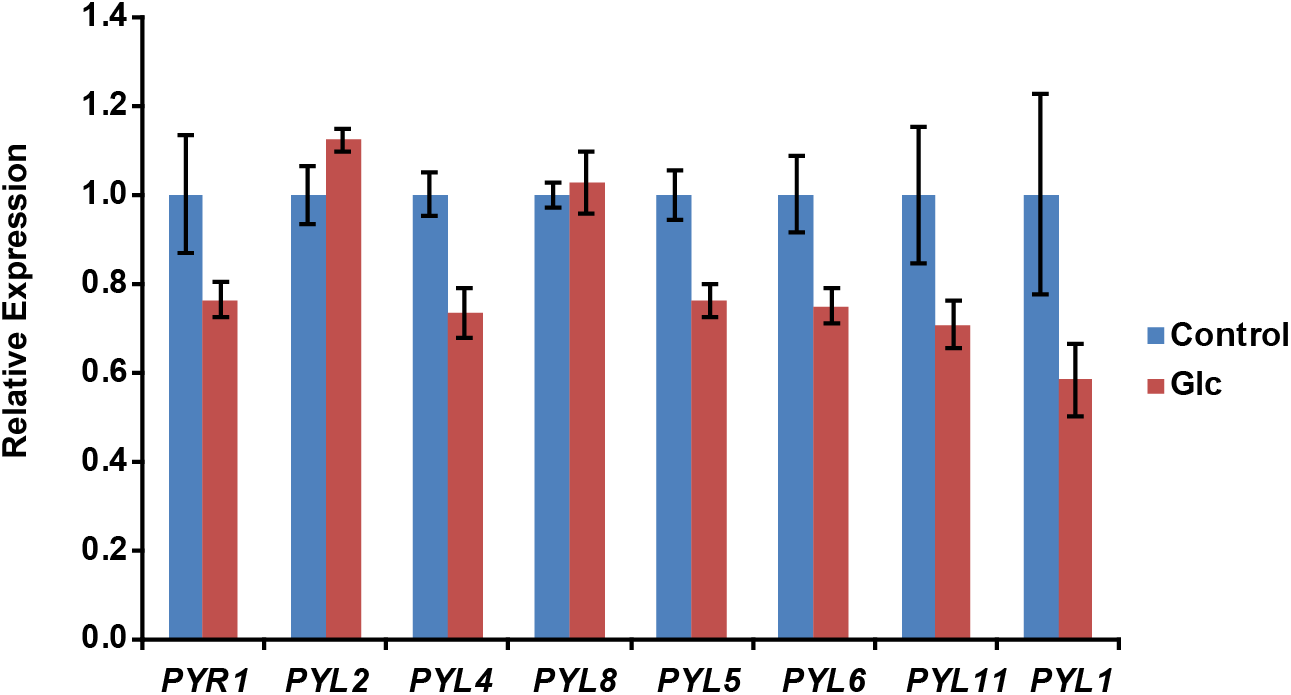
Expression of ABA receptor genes were suppressed by exogenous Glc. Among eight ABA receptor genes (TPM > 10), six were down-regulated by Glc during germination.

**SI Appendix, Table 1.** KEGG pathways enriched in DEG dataset.

**SI Appendix, Table 2.** Gene list and annotation of each cluster.

**SI Appendix, Table 3.** Dormancy associated genes are overrepresented in cluster 1. Among 2127 genes in cluster 1, 637 are associated with dormancy. The list of dormancy associated genes was downloaded from previous study (Bassel et al., 2011)(Bassel *et al*., 2011).

